# *Arabidopsis thaliana* Iron Superoxide Dismutase FeSOD1 Protects ARGONAUTE 1 in a Copper-Dependent Manner

**DOI:** 10.1101/2025.02.26.640378

**Authors:** Ariel H. Tomassi, Ana Perea-García, Guillermo Rodrigo, Javier Sánchez-Vicente, Adriana E. Cisneros, Marta Olmos, Amparo Andrés-Bordería, Lucio López-Dolz, José- Antonio Daròs, Lola Peñarrubia, Alberto Carbonell

**Author notes:** Contributed equally (Ariel H. Tomassi, Ana Perea-García).

## Abstract

Copper (Cu) deficiency compromises plant growth and limits crop productivity. Plants respond to Cu scarcity by activating the expression of several microRNAs, known as Cu-miRNAs, which degrade mRNAs from various cuproproteins to conserve Cu. Cu-miRNAs, like most plant miRNAs, associate with ARGONAUTE 1 (AGO1), the primary effector protein of miRNA-mediated gene silencing pathways, whose function is typically modulated by interacting proteins acting as cofactors. However, how AGO1 is regulated and functions under Cu deficiency remains unknown. Here, we searched for AGO1 interactors in *Arabidopsis thaliana* plants expressing a functional AGO1 protein tagged with the Twin-Strep-tag (TST) polypeptide, grown under Cu-sufficient or Cu-deficient conditions. TST-AGO1 complexes were affinity-purified, and proteins were identified using tandem mass spectrometry. Interestingly, the iron superoxide dismutase 1 (FeSOD1) encoded by *FSD1*, was enriched in TST-AGO1 complexes purified from plants grown under Cu deficiency. Moreover, *fsd1-2* mutant plants showed reduced levels of AGO1 compared to wild-type plants under Cu sufficiency, while both Cu-miRNA-specific and general AGO1 target mRNAs accumulated to higher levels in *fsd1-2* plants under both Cu-deficient and Cu-sufficient conditions compared to wild-type plants. These findings suggest that FeSOD1 is essential for proper AGO1 function, and that its superoxide dismutase activity, which mitigates oxidative stress, enhances AGO1 stability, particularly under Cu deficiency.

**HIGHLIGHT:** AGO1 is essential for Cu-deficiency responses but is sensitive to oxidative stress. FeSOD1 interacts with AGO1 and protects it from superoxide radical-induced degradation, thereby preserving miRNA-mediated gene silencing pathways.

## INTRODUCTION

Plants require rapid and flexible responses to changing environmental conditions throughout their developmental stages, particularly under stress. MicroRNAs (miRNAs) are a class of small RNAs (sRNAs) that regulate gene expression post-transcriptionally and function as key players in the interplay between development and stress responses (Borges and Martienssen, 2015; Song *et al*., 2019). miRNAs regulate a wide range of developmental processes including hypocotyl elongation, root development and the transition from vegetative to reproductive phases (Schwab *et al*., 2005; Meng *et al*., 2010). Conversely, certain miRNAs are integral to stress responses, modulating plant adaptation to a multitude of environmental conditions such as nutrient availability, low temperatures and drought (Patel *et al*., 2017; Martin *et al*., 2010; Buhtz *et al*., 2010). For instance, miRNAs involved in nutrient stress, including transition metal deficiencies, play significant roles in plant adaptation (Patel *et al*., 2017) by regulating processes like metal uptake, chelation, antioxidant response and hormone signaling (Ding *et al*., 2020).

The availability of essential transition metals, such as copper (Cu) and iron (Fe), often limit agricultural productivity (Broadley *et al*., 2012). These metals, essentials as cofactors in metalloproteins central to life, can also cause toxicity when in excess, leading to increased reactive oxidative species (ROS) production (Farooq *et al*., 2019). Due to their dual nature as essential and toxic elements, plants have developed complex homeostatic networks to assure their uptake and distribution while preventing toxicity (Puig and Peñarrubia, 2009; Pilon, 2011; Kobayashi *et al*., 2019; Kumar *et al*., 2021). ROS are also produced under metal scarcity, as the malfunctioning of key redox metalloproteins in respiratory and photosynthetic electron transport chains causes oxidative stress (Shahbaz and Pilon, 2019). Among ROS, the superoxide radical is one of the most harmful species, capable of damaging macromolecules. ROS detoxification involves superoxide dismutases (SODs), which in plants use different metals as cofactors, functioning in various cellular compartments under diverse conditions. Plant SODs include FeSODs, Cu and zinc (Zn)SODs (Cu/ZnSODs) and manganese (Mn) MnSODs (Ravet and Pilon, 2013).

In *Arabidopsis thaliana* (Arabidopsis), the denoted as Cu-miRNAs play crucial roles in plant responses under Cu deficiency, orchestrating metal redistribution and substituting Cu/ZnSODs with FeSODs (Yamasaki *et al*., 2007; Pilon, 2017). Cu-miRNAs are proposed to modulate metalloprotein mRNAs post-transcriptionally, establishing a metal prioritization hierarchy and preventing interference with metal-sensitive processes (Perea-García *et al*., 2022; Shahbaz and Pilon, 2019). Cu-miRNAs, including miR397, miR398, miR408 and miR857, are induced under Cu deficiency and primarily target mRNAs encoding Cu/ZnSODs (*CSD1* and *CSD2*), plantacyanin and laccases (Pilon, 2017). Cu-miRNAs containing *cis*- regulatory domains in their promoters, recognized by a SQUAMOSA BINDING PROTEIN-LIKE (SPL) family member, SPL7, the master transcription factor driving Cu-deficiency responses (Yamasaki *et al*., 2009; Bernal *et al*., 2012). In addition to Cu-miRNAs, plasma membrane Cu transporters genes (*COPT1*, *COPT2* and *COPT6*) as well as *FSD1* (encoding FeSOD1) are regulated similarly (Yamasaki *et al*., 2007; Bernal *et al*., 2012; Garcia-Molina *et al*., 2013). These processes enhance high affinity Cu uptake and metalloprotein substitution as primary responses to nutrient scarcity.

Importantly, SPL7 belongs to a large family of transcription factors mainly involved in the control of developmental phase transitions and reproduction (Huijser and Schmid, 2011). Based on their sizes and similarities, the Arabidopsis SPL family can be grouped in two subfamilies: large SPLs (including SPL1, SPL7, SPL12, SPL14 and SPL16) and small SPLs (the remaining 11 SPLs) (Guo *et al*., 2008; Xing *et al*., 2010). The small SPLs, with the exception of SPL8, share a miRNA response element that is complementary to miR156 responsive in the timing of the vegetative and reproductive phase transitions (Rhoades *et al*., 2002; Schwab *et al*., 2005; Wu and Poethig, 2006; Gandikota *et al*., 2007). *SPL3* overexpressing plants, resistant to miR156 regulation, display a strong reduction in SPL7- mediated Cu-deficiency responses, probably due to competition between SPL7 and miR156- regulated SPL3 for *cis* element binding in target promoters (Perea-García *et al*., 2021). Therefore, the balance of miR156 SPL targets and non-targets is essential for SPL7-mediated Cu-deficiency adaptation at different levels, underscoring that the effect of miRNA on its targets could serve to reinforce the miRNA non-target function. Thus, while Cu-miRNAs function as post-transcriptional modulators of metalloproteins, allowing flexible Cu redistribution, other miRNAs, such as miR156, operate upstream, balancing the relative abundance of SPL transcription factors to integrate Cu-deficiency responses with developmental processes.

ARGONAUTE (AGO) proteins act as the effector proteins in sRNA-mediated RNA silencing pathways in eukaryotes (Meister, 2013). In plants, AGOs regulate essential biological processes such as development, stress responses, genome stability and defense against pathogens, particularly to viruses (Carbonell, 2017; Fang and Qi, 2016; Carbonell and Carrington, 2015). They associate with sRNAs such as miRNAs and siRNAs to form the RNA-induced silencing complex (RISC), which directs the endonucleolytic cleavage or translational repression of sequence-complementary target RNAs (Huntzinger and Izaurralde, 2011).

Arabidopsis has 10 AGO members that have functionally diverged during the specialization of sRNA-based RNA silencing pathways (Chapman and Carrington, 2007). AGO1 is the principal miRNA-associating AGO (Vaucheret *et al*., 2004) and, consequently, *ago1* mutants show severe growth defects (Morel *et al*., 2002). AGO1 protein levels, are regulated at different transcriptional and posttranscriptional levels (Molinier, 2020). For instance, AGO1 can be ubiquitinated and degraded through the ubiquitin-proteasome pathway that tags AGOs with ubiquitin molecules for their degradation by the 26S proteasome (Earley *et al*., 2010), or through autophagy by sequestration of AGO1 proteins in autophagosomes (Derrien *et al*., 2012). In contrast, AGO1 protein is stabilized by its binding to sRNAs (Vaucheret *et al*., 2006; Hacquard *et al*., 2022), as unloaded AGO1 is more prone to degradation, by its sequestration into stress granules (Leung *et al*., 2006) or through its association with co-factors. AGO1 physically interacts with several proteins, including chaperones involved in RISC loading such as HEAT SHOCK PROTEIN 70 (HSP70), HSP90 and CYCLOPHILLIN 40/SQUINT (CYP40/SQN), which facilitate the conformational opening of AGO1 during sRNA loading (Iki *et al*., 2010; Iwasaki *et al*., 2010; Iki *et al*., 2012). Moreover, TRANSPORTIN 1 (TRN1), ENHANCED MiRNA ACTIVITY 1 (EMA1) β and CONSTITUTIVE ALTERATIONS IN THE SMALL RNAS PATHWAYS 9 (CARP9) proteins interact with AGO1 to modulate miRNA loading (Wang *et al*., 2011; Cui *et al*., 2016; Tomassi *et al*., 2020), while PROTEIN ARGININE METHYLTRANSFERASE 5 (PRMT5) associate with AGO1 to promote its methylation (Martín-Merchán *et al*., 2024). Finally, proteins known to interact with AGO1 in response to stress are limited to DAMAGE-SPECIFIC DNA BINDING PROTEIN 2 (DDB2), TUDOR DOMAN-CONTAINING PROTEIN 1 (TSN1) and TSN2 and SWI/SNF complexes in response to UV, general stress and hormone or cold stress, respectively (Liu *et al*., 2018; Gutierrez-Beltran *et al*., 2021; Schalk *et al*., 2017). However, proteins specifically interacting with AGO1 during the metal stress response have not been identified.

To better understand AGO1’s role in response to Cu deficiency, we searched for AGO1 interactors in Arabidopsis transgenic lines expressing a functional AGO1 protein tagged with the bidentate polypeptide Twin-Strep-tag (TST), grown under Cu-sufficient or Cu-deficient conditions. AGO1 complexes were affinity-purified, and proteins were identified by tandem mass spectrometry (AP-MS or affinity purification mass spectrometry). Interestingly, *FSD1*- encoded FeSOD1, recently proposed as an AGO1 interactor as a result of a yeast two-hybrid screening (Bressendorff *et al*., 2023), was enriched in AGO1 complexes purified from plants grown under Cu deficiency. *fsd1-2* mutant plants accumulated lower levels of AGO1 compared to Col-0 plants under Cu sufficiency, and both Cu-miRNA-specific and general AGO1 target mRNAs accumulated to higher levels in *fsd1-2* plants grown under Cu deficiency and sufficiency, respectively, compared to Col-0. These results underscore the role of FeSOD1 in protecting AGO1 stability, especially under Cu deficiency, where its function in miRNA-mediated gene silencing pathways becomes particularly important.

## RESULTS

### Generation and functional characterization of Arabidopsis transgenic lines expressing TST-tagged AGO1

TST is an advanced affinity tag used for the purification and detection of recombinant proteins under non-denaturing, mild conditions (Schmidt *et al*., 2013). It consists of a tandem arrangement of two Strep-tag II sequences, designed to enhance binding affinity to Strep-Tactin, a derivative of streptavidin. The TST coding sequence was inserted after AGO1 start codon to tag the N-terminus of AGO1, a previously done with hemagglutinin (HA) or FLAG peptides (Baumberger and Baulcombe, 2005; Carbonell *et al*., 2012), in the *pAGO1:TST-AGO1* construct, which includes authentic AGO1 regulatory sequences (Figure 1A). As a control, the *pAGO1:TST-GFP* construct, designed to express TST-tagged GFP under the same AGO1 regulatory sequences, was also generated (Figure 1A). To test the functionality of TST-AGO1, *pAGO1:TST-AGO1* and *pAGO1:TST-GFP* constructs were transformed in both Col-0 and *ago1-25* genetic backgrounds. At least, 24 independent T1 transgenic lines were recovered for each construct in both backgrounds. Interestingly, *ago1-25* phenotypic defects, such as slow growth or delayed flowering, were suppressed in plants expressing *pAGO1:TST-AGO1*, but not in plants expressing *pAGO1:TST-GFP* (Figure 1B and 1C). In contrast, plants expressing either construct in the Col-0 background were phenotypically similar to non-transformed Col-0 plants (Figure 1B and 1C). Importantly, plants expressing *pAGO1:TST-AGO1* or *pAGO1-TST-GFP* constructs accumulated high levels of TST-AGO1 and TST-GFP proteins, respectively, as observed in the western blot analysis (Figure 1D). To further investigate TST-AGO1 functionality, AGO1-dependent *TAS1c* tasiRNA 255 (tasiR255) formation was analyzed in T1 transgenic plants. TasiR255 accumulation levels were restored to Col-0 levels in *ago1-25* plants transformed with *pAGO1:TST-AGO1* but not in *ago1-25* plants transformed with *pAGO1:TST-GFP* (Figure 1D). Finally, tasiR255 levels in transgenic lines expressing either construct in the Col-0 background were similar than those of non-transformed Col-0 plants. Overall, these results indicate that the addition of a TST tag at the 5’ terminus of Arabidopsis AGO1 does not affect AGO1 functionality. They also suggest that the expression of TST-AGO1 in a Col-0 background does not impair plant growth or AGO1 functionality.

**Figure 1.**
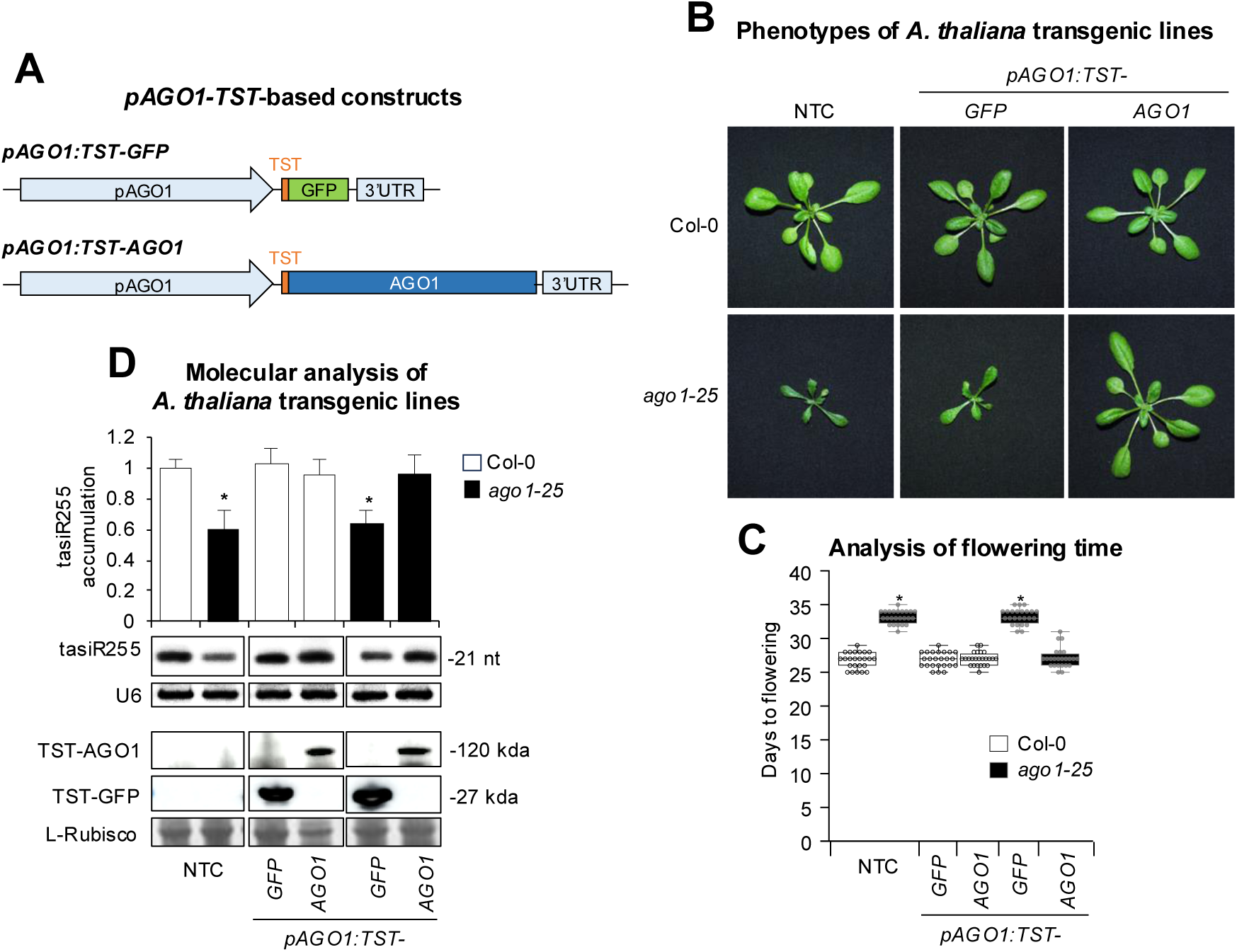
Phenotypic and molecular analyses of Arabidopsis Col-0 and *ago1-25* transgenic plants expressing TST-tagged AGO1 or GFP. **A**, Diagram of the constructs. TST, GFP and AGO1 sequences are shown in orange, green and dark blue boxes, respectively. AGO1 endogenous promoter and 3’ UTR regulatory sequences are shown in a light blue arrow and box, respectively. **B**, Pictures of 21-day-old Col-0 (top panel) and *ago1-25* (bottom panel) T1 transgenic plants, homozygous for the transgene. Col-0 and *ago1-25* non-transgenic controls (NTCs) are included. **C**, Box plot representing the mean flowering time of Col-0 and *ago1-25* NTC and T1 transgenic plants (*n* = 24). Pairwise Student’s *t*-test comparisons are represented with an * if significantly different (*P* < 0.05) compared to Col-0 NTC. **D**, Accumulation of *TAS1c*-dependent tasiRNA (tasiR255) and TST-AGO1 in T1 transgenic lines. The graph at the top shows the mean (*n* = 3) relative to tasiR255 levels + standard deviation (Col-0 = 1.0). One blot from three biological replicates is shown. Each biological replicate is a pool of at least eight independent lines that were randomly selected. U6 and L-Rubisco (ribulose-1,5-bisphosphate carboxylase/oxygenase) blots are shown as loading controls. Other details are as in C.

### AP-MS analysis of AGO1 protein complexes in response to Cu deficiency stress

Given the previously described key role of miRNA-mediate gene regulation in response to Cu deficiency (Pilon, 2017; Perea-García *et al*., 2022), we analyzed the AGO1 protein interactome under Cu deficiency compared to Cu sufficiency, using three independent T4 pAGO1:TST-AGO1 transgenic lines to study AGO1 function in both conditions: lines 1, 2 and 3. As a negative control, one T4 *pAGO1:TST-GFP* transgenic line was also grown in parallel under the same conditions. Seedlings of each transgenic line were grown for 10 days under Cu sufficiency (1 μM CuSO4) or under Cu deficiency (100 μM bathocuproine disulphonate, BCS) conditions. AGO1 and GFP protein complexes were purified by affinity chromatography in native conditions. Proteins from the purified complexes were resolved using denaturing polyacrylamide gel electrophoresis (PAGE) and subsequently subjected to in-gel digestion with trypsin. The resulting peptides were extracted from the gel and analyzed by liquid chromatography coupled with tandem mass spectrometry to identify the proteins in the purified complexes. Data S1 shows all hits resulting from proteomic analysis of complexes purified from seedlings of each transgenic line grown under Cu sufficiency and Cu deficiency.

Next, we examined the dynamics of the AGO1 protein interactome in response to Cu deficiency. Proteins from purified samples were processed using the ProteinPilot (Shilov *et al*., 2007) and Mascot (Perkins *et al*., 1999) software tools, and the identified proteins were matched against the *Viridiplantae* protein database to filter for plant-specific proteins (Figure 2A). These plant proteins were further analyzed using the BLAST tool (Altschul *et al*., 1990) against the NCBI Genomes to identify Arabidopsis-specific proteins. The total number of proteins identified in the three *pAGO1:TST-AGO1* lines under Cu deficiency were similar (2185, 2099 and 2002 for lines 1, 2 and 3, respectively), although higher than the number observed in all lines under Cu sufficiency (1565, 1680 and 1749 for lines 1, 2 and 3, respectively) (Figure 2B). The total number of proteins identified in the *pAGO1:TST-GFP* line under Cu deficiency and sufficiency was rather similar (2080 and 1975, respectively) (Figure 2B).We then quantified the change in the abundance of AP-MS interactors under these two conditions (Data S2), discarding those proteins present in the *pAGO1:TST-GFP* control purifications. A log2 fold change value was assigned for each protein comparison, with log2 fold > 0 indicating that the protein is more abundant in Cu deficiency and log2 fold < 0 indicating higher abundance in Cu sufficiency, respectively. A Venn diagram showed that 89 proteins were more abundant in Cu deficiency than in Cu sufficiency in the three lines (Figure 2C, Data S3). Among these proteins, we considered the Fe superoxide dismutase 1 protein (FeSOD1) for our subsequent analyses, as it was recently identified as an AGO1 interactor in yeast two hybrid assays (Bressendorff *et al*., 2023) and was also known to participate in oxidative stress responses in a Cu-dependent manner (Melicher *et al*., 2022).

**Figure 2.**
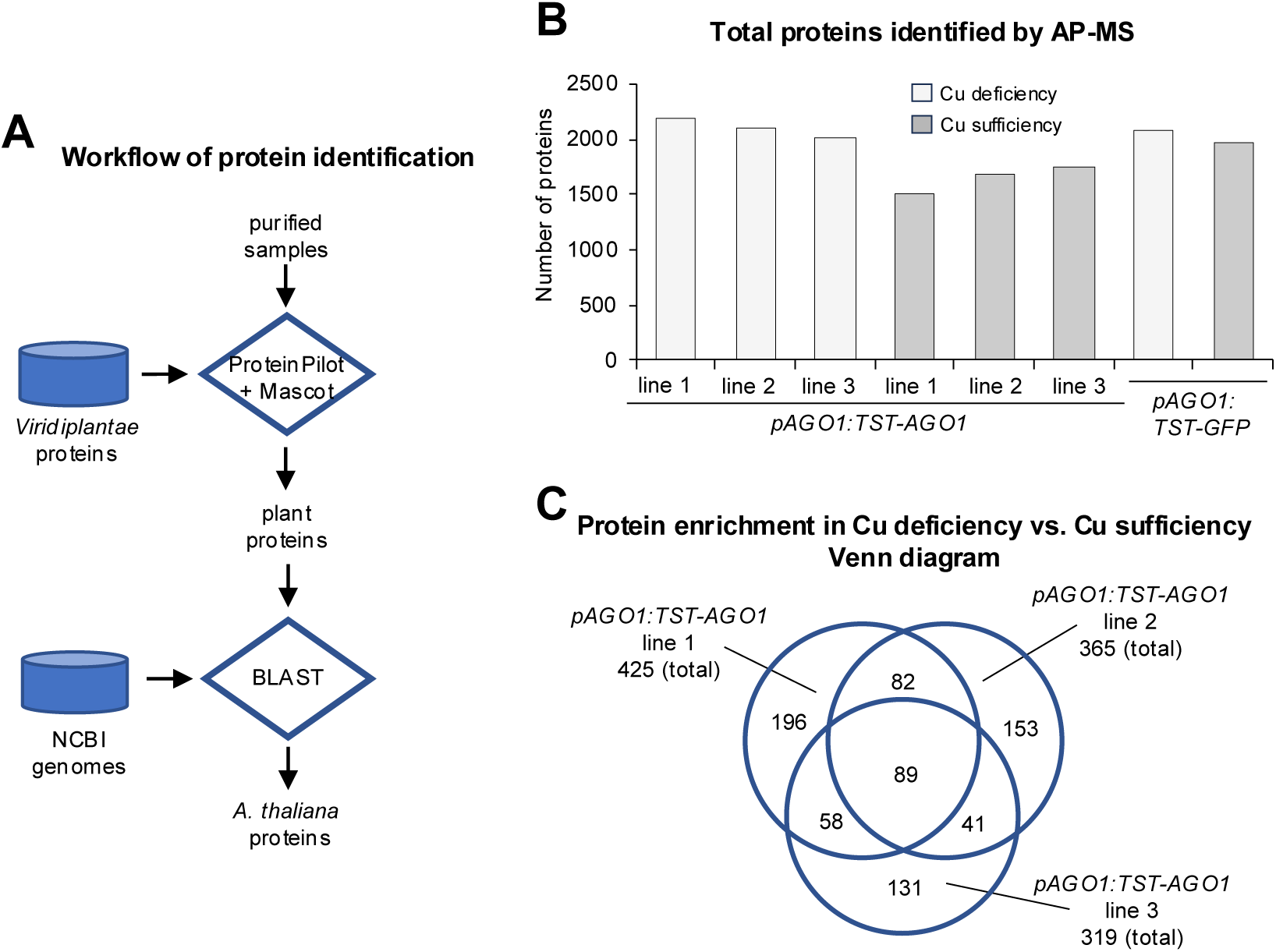
**Protein analyses in TST-AGO1 complexes purified by AP-MS**. **A**, Flowchart of the process for plant protein identification. Proteins from purified samples are processed through ProteinPilot and Mascot software tools, and identified proteins are matched against the *Viridiplantae* protein database to filter for plant proteins. The plant proteins are further analyzed using the Blast tool against the NCBI Genomes database to identify Arabidopsis-specific proteins. **B**, Bar chart showing the total number of proteins identified by AP-MS in the three different *pAGO1-TST-AGO1* transgenic lines grown under copper (Cu) deficiency (light grey bars) or Cu sufficiency (dark grey bars). **C**, Venn diagram comparing the total number of proteins in Cu deficiency versus Cu sufficiency identified in the three *pAGO1:TST-AGO1* transgenic lines. The cutoff value for significant fold change is set at log2 fold = 1, meaning that only proteins enriched in Cu deficiency versus Cu sufficiency are represented. Overlapping section indicates the proteins identified in purified samples from independent transgenic lines.

### AGO1 and FeSOD1 fine-tune Cu-responsive SOD gene expression

To assess the impact of Cu deficiency on plant growth, Arabidopsis Col-0, *fsd1-2* (loss-of-function mutant for *FSD1*), and *ago1-25* (mutant for *AGO1*) seedlings were grown under Cu-sufficient and Cu-deficient conditions (Figure 3A), and their root lengths were quantified (Figure 3B). Under Cu sufficiency, all genotypes displayed healthy growth, with *fsd1-2* seedlings showing a slight but statistically significant increase in root length compared to Col-0 (*P* < 0.05), while *ago1-25* seedlings exhibiting a pronounced decrease as observed before in other *ago1* mutants (Trolet *et al*., 2019). Under Cu deficiency, root growth was significantly inhibited across all genotypes (Figure 3A, 3B).

**Figure 3.**
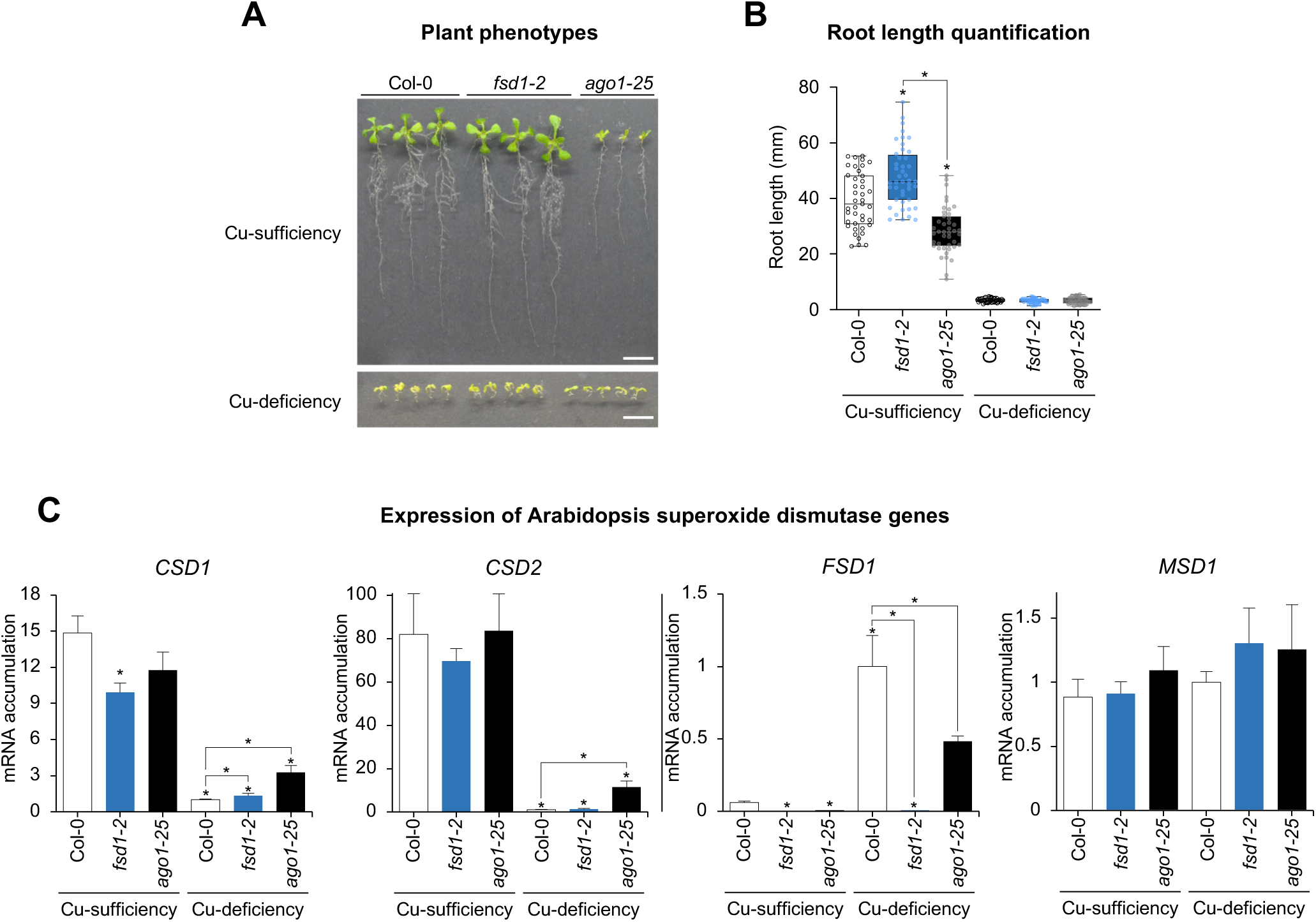
Analysis of Arabidopsis superoxide dismutase genes under copper sufficiency and deficiency conditions in Col-0, *fsd1-2* and *ago1-25* seedlings. **A**, Photographs of 10-day-old of Col-0, *fsd1-2* and *ago1-25* seedlings grown on plates with MS medium in Cu sufficiency (1 µM Cu) or deficiency (100 µM BCS). **B**, Quantification of root length of Col-0, *fsd1-2* and *ago1-25* seedlings grown in the same conditions than in A. Mean relative level (*n* = 40-45) + standard error of root length. Pairwise Student’s *t*-test comparisons are represented with an * if significantly different (*P* < 0.05) compared to Col-0 in each condition. **C**, Relative expression of *CSD1, CSD2, FSD1* and *MSD1* mRNAs determined by RT-qPCR in 10-day-old Col-0, *fsd1-2* and *ago1-25* seedlings, grown under control (Cu sufficiency, 1 μM CuSO4) and Cu deficiency (100 μM BCS). Mean relative level (*n* = 3) + standard error of mRNAs after normalization to *ACTIN2*. Bars with an * are significantly different from that of the Col-0 control sample (*P* < 0.05 in all pairwise Student’s *t*-test comparisons). Other *t*-test comparisons are also shown. Each biological replicate is a pool of at least twenty independent lines that were randomly selected.

Expression of SOD genes is an excellent marker for Cu status in plant cells (Yamasaki *et al*., 2009; Andrés-Colás *et al*., 2013). To examine SOD expression under our experimental conditions, mRNA accumulation of *CSD1*, *CSD2*, *FSD1* and *MSD1* was measured by RT-qPCR (Figure 3C) in Col-0, *fsd1-2* and *ago1-25* seedlings. As previously reported (Yamasaki *et al*., 2009), expression of Cu/Zn SODs encoded by *CSD1* and *CSD2* in Col-0 seedlings was high under Cu sufficiency but remained low under Cu deficiency (Figure 3A). Although the expression pattern in *ago1-25* was similar, *CSD* expression was significantly higher in most cases (Figure 3C). Since *CSD* expression is regulated by *miR398*, the impaired AGO1 function in *ago1-25* could explain this difference. On the other hand, expression of FeSOD1, encoded by *FSD1,* remained low under Cu sufficiency and was higher under Cu deficiency in both genotypes, although it was significantly lower in *ago1-25* under Cu deficiency (Figure 3C). This suggests an indirect effect of AGO1 function on *FSD1* transcription under Cu deficiency, potentially due to miR156 influencing the ratio of different SPL members and thereby affecting SPL7’s occupancy of Cu deficiency target promoters (Perea-García *et al*., 2021). Finally, expression of MnSOD, encoded by MSD1, remained approximately constant across both genotypes and conditions (Figure 3C).

### FeSOD1 is essential for AGO1 stability and functionality, particularly under Cu deficiency

Next, we investigated whether FeSOD1 function was required for proper AGO1 expression. To do so, AGO1 protein levels were measured by western blot in 10-day-old Col-0, *fsd1-2* and *ago1-25* seedlings grown under Cu sufficiency and Cu deficiency conditions (Figure 4A). Surprisingly, AGO1 protein levels in Col-0 plants drastically dropped (around 90%) in Cu deficiency compared to Cu sufficiency. This decrease was similar to that observed in *ago1-25* plants grown under both Cu conditions. Additionally, AGO1 protein levels also decreased by about 80% in *fsd1-2* seedlings compared to Col-0 plants under Cu sufficiency (Figure 4A). These results indicate an important decrease in AGO1 protein in Col-0 plants grown content under Cu deficiency, and in *fsd1-2* seedlings grown under both Cu conditions.

**Figure 4.**
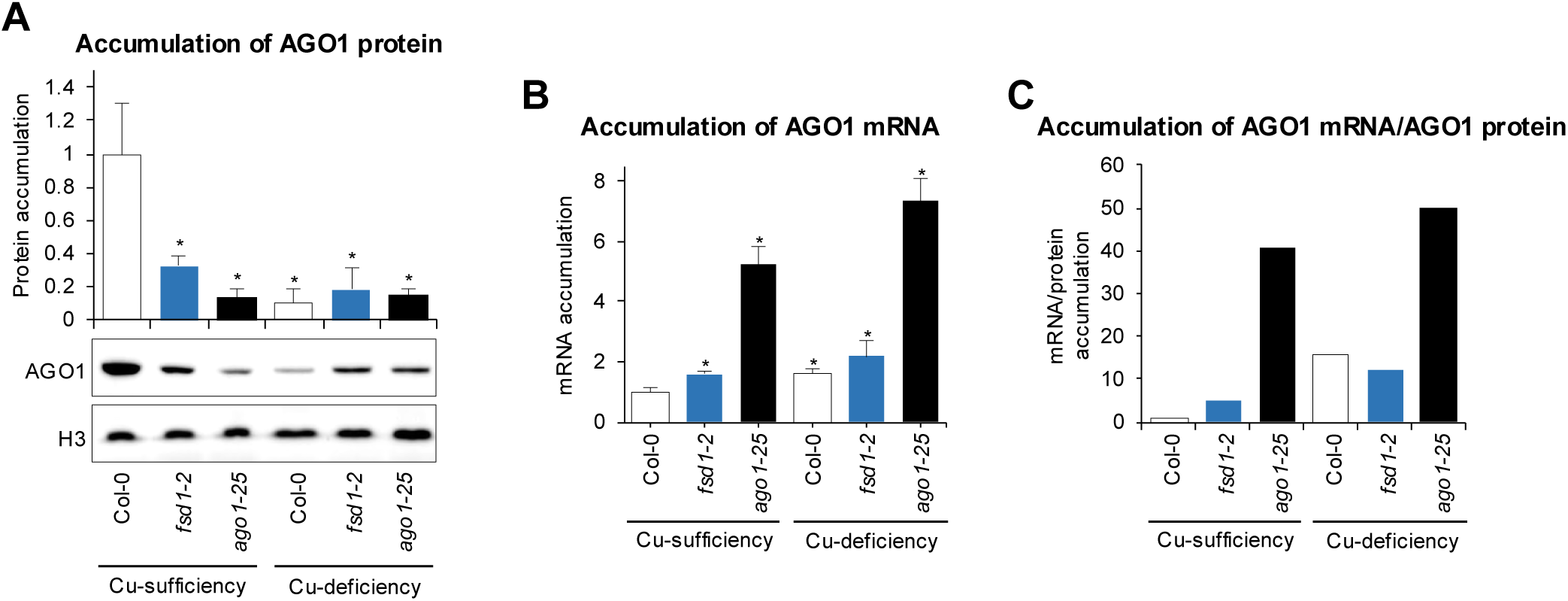
Accumulation of AGO1 protein and mRNA in Arabidopsis Col-0*, fsd1-2* and *ago1-25* seedlings. **A**, Accumulation of endogenous AGO1 protein in 10-day-old Col-0, *fsd1-2* and *ago1-25* seedlings, grown under control (Cu sufficiency) and Cu deficiency. The graph at the top shows the mean (*n* = 3) relative to endogenous AGO1 levels + standard deviation (Col-0 = 1.0). One blot from three biological replicates is shown. Each biological replicate is a pool of at least twenty independent lines that were randomly selected. HISTONE H3 (H3) blot is shown as loading control. **B**, Relative expression of *AGO1* determined by RT-qPCR in 10-day-old Col-0, *fsd1-2* and *ago1-25* seedlings, grown under control and Cu deficiency. Mean relative level (*n* = 3) + standard error of mRNA after normalization to *ACTIN2* (Col-0 = 1.0). Bars with an * are significantly different from that of the Col-0 control sample (*P* < 0.05 in all pairwise Student’s *t*-test comparisons). Each biological replicate is a pool of at least twenty independent lines that were randomly selected.

To further investigate whether the reduction in AGO1 protein levels was due to transcriptional regulation, we analyzed *AGO1* mRNA levels by RT-qPCR (Figure 4B) in the same samples used for AGO1 protein analysis (Figure 4A). Under Cu sufficiency, *AGO1* transcript levels were significantly higher in *fsd1-2* compared to Col-0, suggesting that FeSOD1 function may influence *AGO1* expression. Additionally, *ago1-25* mutants exhibited a significant increase in *AGO1* mRNA levels relative to Col-0, likely due to impaired miR168-mediated AGO1 mRNA regulation caused by reduced AGO1 activity (Vaucheret *et al*., 2006). Under Cu deficiency, *AGO1* transcript accumulation markedly increased in Col-0, *fsd1-2* and *ago1-25* seedlings compared to Cu sufficiency, with *ago1-25* displaying the highest expression levels (Figure 4B). These results indicate that the reduction in AGO1 protein levels under Cu deficiency is not due to decreased AGO1 transcript accumulation.

To further assess the relationship between *AGO1* mRNA and protein levels, we calculated the relative ratio of *AGO1* mRNA to AGO1 protein accumulation in the same samples (Figure 4C). Under Cu sufficiency, this ratio (R) was much higher in *fsd1-2* (R=4.9) and *ago1-25* (R=40.6) mutants compared to Col-0 (R=1), indicating that despite increased *AGO1* transcript levels, AGO1 protein accumulation remained low in these mutants. Under Cu deficiency, the *AGO1* mRNA/protein ratio increased more drastically across all genotypes, particularly in *ago1-25*, which exhibited the highest ratio (R=50.2) among the conditions tested.

To determine whether the decrease in AGO1 protein content also affected its functionality under Cu deficiency, the expression of several Cu-miRNA targets (*ARPN,* miR160 target; *CCS, CSD1, CSD2,* miR398 targets; GST25, miR396 target; *LAC3*, miR397 target) was analyzed by RT-qPCR in 10-day-old Col-0, *fsd1-2* and *ago1-25* seedlings (Figure 5A). As expected, the expression of all Cu-miRNA targets (except for *GSTU25*) significantly increased in *ago1-25* compared to Col-0. Similarly, all miRNA targets were induced to varying degrees in *fsd1-2* compared to Col-0, with the increase in *CSD1* being statistically significant (Figure 5A). These findings confirm that AGO1 functionality requires the presence of FeSOD1 under Cu deficiency.

**Figure 5.**
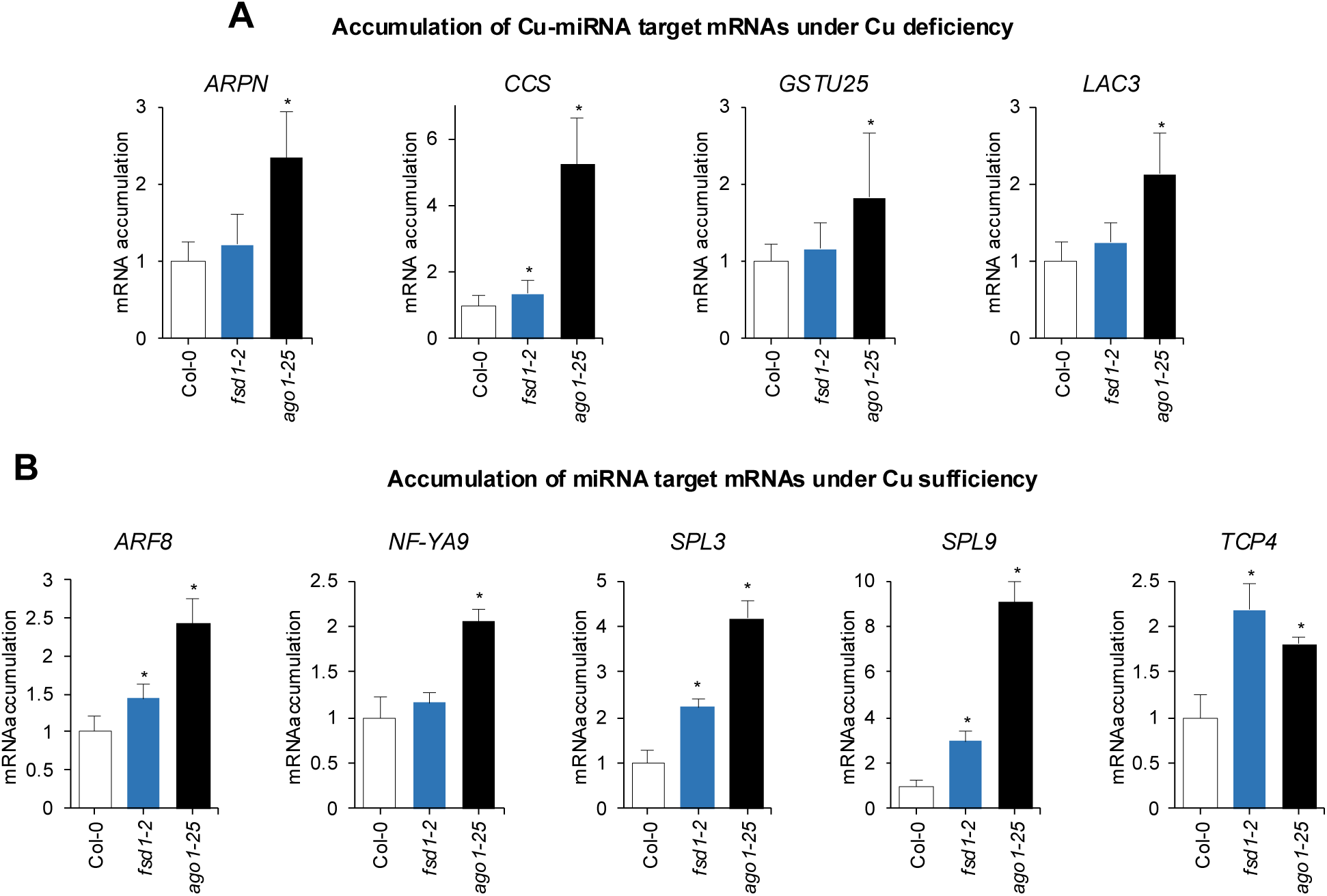
Accumulation of miRNA target genes in Arabidopsis Col-0*, fsd1-2* and *ago1-25* seedlings. **A**, Relative expression of Cu-miRNA targets *ARPN*, *CCS*, *LAC3* and *GSTU25* determined by RT-qPCR in 10-day-old Col-0, *fsd1-2* and *ago1-25* seedlings, grown under Cu deficiency. Mean relative level (*n* = 3) + standard error of mRNAs after normalization to *ACTIN2* (Col-0 = 1.0). Bars with an * are significantly different from that of the Col-0 control sample (*P* < 0.05 in all pairwise Student’s *t*-test comparisons). Each biological replicate is a pool of at least twenty independent lines that were randomly selected. **B,** Relative expression of miRNA targets *ARF8*, *NF-YA9, SPL3*, *SPL9* and *TCP4* determined by RT-qPCR in 10-day-old Col-0, *fsd1-2* and *ago1-25* seedlings, grown under Cu sufficiency (1 μM CuSO4). Other details are as in A.

Next, we wondered whether FeSOD1 activity was also required for general AGO1 function under Cu sufficiency and for regulating other non-Cu-miRNA targets. The expression of different well-known miRNA targets (*ARF8*, miR167 target; *NF-YA9*, miR169 target; *SPL3* and *SPL9*, miR156 targets; *TCP4*, miR159 target) was analyzed by RT-qPCR in 10-day-old Col-0, *fsd1-2* and *ago1-25* seedlings grown under Cu sufficiency (Figure 5B). As expected, the expression of all miRNA targets was significantly higher in *ago1-25* compared to Col-0. Interestingly, the expression of all miRNA targets also increased in *fsd1-2* compared to Col-0, with a statistically significant increase observed for four out of five miRNA targets (*ARF8*, *SPL3*, *SPL9* and *TCP4*). Notably, the increased expression of miR156 targets (*SPL3* and *SPL9*) is relevant, as these targets compete with SPL7 in Cu-deficiency responses (Perea-García *et al*., 2021). Finally, the expression of the miRNA targets mentioned above was also analyzed in seedlings of the different genotypes grown under Cu deficiency (Figure S1). The expression of all miRNA targets was higher in both *fsd1-2* and *ago1-25* compared to Col-0, although this increase was significant only for *NF-YA9* and *SPL3* in *ago1-25* (Figure S1).

In conclusion, our results indicate that FeSOD1 is necessary for proper AGO1 function in miRNA-mediated gene silencing. We propose a model (Figure 6) in which FeSOD1’s superoxide dismutase activity protects against oxidative stress, contributing to the increased stability of the AGO1 protein. This protective role of FeSOD1 is especially important under Cu deficiency due to the elevated oxidative stress and the critical role played by Cu-miRNAs in the plant’s response to this nutrient deficit. Absence of FeSOD1 in *fsd1-2* mutants results in AGO1 degradation independently of the Cu levels (Figure 6).

**Figure 6.**
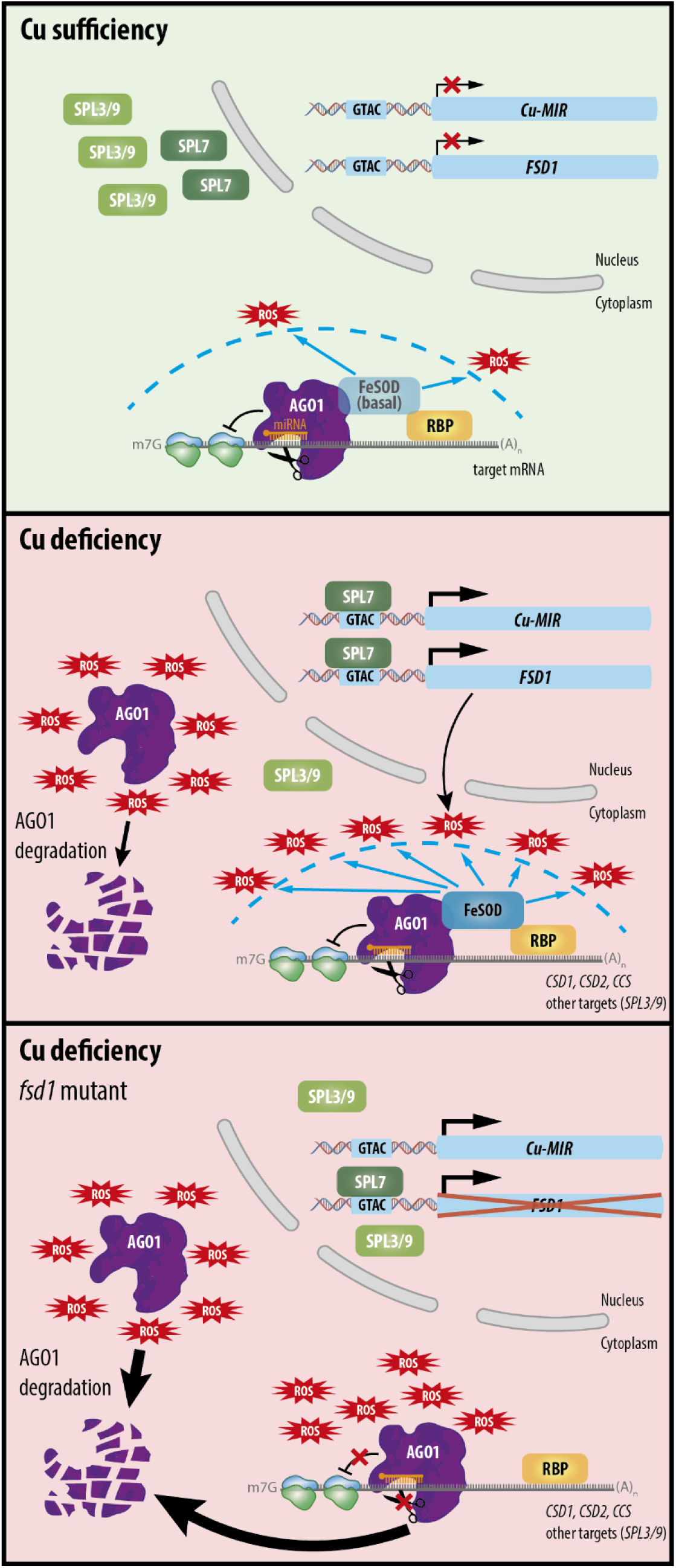
Model for the anti-superoxide protection of the AGO1 function by the FeSOD1 dome under Cu deficiency. Top, under Cu sufficiency Cu-miRNA and *FSD1* expression are essentially blocked. A basal level of iron superoxide dismutase FeSOD1 allows the protection of AGO1 from reactive oxygen species (ROS) derived from normal cellular metabolism. Middle, under Cu deficiency, the increase presence of ROS, including superoxide anion, induces a certain level of AGO1 degradation. However, in these conditions SPL7 outcompetes SPL3 and SPL9 for binding to cis-regulatory transcriptional elements (GTACs) present in promoters of Cu-miRNA genes and FeSOD1-encoding *FSD1,* inducing their expression. Increased levels of FeSOD1 play a protective role of the effector complex of RNAi by detoxifying superoxide anion. By directly interacting with AGO1 and/or associated RNA Binding Proteins (RBP), FeSOD1 promotes AGO1 stability contributing to Cu-miRNA and general AGO1 functions. Bottom, in *fsd1* mutants, the lack of FeSOD impedes proper detoxification of ROS and leads to AGO1 degradation in both Cu sufficiency (left) and Cu deficiency (right) conditions.

## DISCUSSION

This study investigates plant responses under Cu deficiency conditions, where Cu-miRNAs regulate the mRNAs encoding cuproproteins. Different studies have reported that numerous miRNAs are induced by a variety of environmental stresses, suggesting the existence of shared signaling pathways (Martin *et al*., 2010). Plants face a conflictive situation when responding to adverse conditions, give the extensive role of miRNAs in stress responses and the instability of the key protein AGO1 within RISC under suboptimal conditions (Fang and Qi, 2016). To solve this conflict, providing a protective environment to preserve AGO1 function could be crucial for an effective stress response. Here, we explored how Cu deficiency influences the regulation of miRNA targets mediated by AGO1. We analyzed the AGO1 protein interactome in *pAGO1:TST-AGO1* Arabidopsis transgenic plants grown under both normal and Cu-deficient conditions. Due to the adverse effects of Cu deficiency on AGO1 function, we searched for interactor proteins with a protective effect on AGO1. Notably, a well-documented response to Cu deficiency is the substitution of Cu/ZnSODs with FeSODs to support antioxidative defense (Yamasaki *et al*., 2009). In this context, FeSOD1 was enriched in TST-AGO1 complexes purified from all three *pAGO1:TST-AGO1* lines grown in Cu deficiency. Since FeSOD1 was recently identified as an AGO1 interactor in yeast two-hybrid assays (Bressendorff *et al*., 2023), the role of FeSOD1-AGO1 interaction became the main focus of our study.

Particularly striking is the opposite pattern observed between *AGO1* mRNA and protein levels, suggesting that either AGO1 translation or stability is severely compromised under Cu deficiency. This is evidenced by the significant decrease in endogenous AGO1 protein levels in Col-0 plants grown under Cu-deficient conditions compared to Cu-sufficient conditions. A similar opposite pattern is also observed in *fsd1-2* and, more drastically, in the *ago1-25* mutant, possibly indicating a compensatory increase in *AGO1* mRNA synthesis in response to low AGO1 protein levels (Figure 4).

The elevated ROS levels induced by Cu deficiency appear to be a key factor explaining AGO1 degradation, as the presence of FeSOD1 detoxifies the cellular oxidative state and safeguards AGO1 function in regulating Cu-miRNA genes or other known targets (Figure 5A, Figure 6, Figure S1). Consistently, as peroxide is the reaction product of SODs, the peroxidase PRX32 is also identified as antioxidant molecules interacting with AGO1 under Cu deficiency (Data S3). Whether ROS-induced AGO1 degradation occurs via the ubiquitin-proteasome system, the autophagy pathway or simultaneously through both remains unknown. It is possible that specific proteins target AGO1 for degradation, as reported for the F-box protein FBW2 that assembles an SCF complex that selectively targets for proteolysis AGO1 when it is unloaded (Hacquard *et al*., 2022).

Alongside Cu-miRNAs, SPL7 can also induce *FSD1* expression (Yamasaki *et al*., 2009). Our results indicate that defects in AGO1 function impact Cu-miRNAs and *FSD1* expression (Figure 3C). The reduced expression of *FSD1* in *ago1-25* likely results from defective miR156 targeting. Indirectly, *FSD1* expression is affected by miR156, which main function is to inhibit the gene expression of some members of the SPL protein family (especially SPL3), while it does not affect other members such as SPL7. The purpose seems to be reducing competition between SPL factors for target promoter binding (Perea-García *et al*., 2021). Consequently, the defective targeting of miR156 in *ago1-25* explains the increased expression of non-SPL7 members (SPL3 and SPL9), potentially disrupting SPL7 responses, including *FSD1* expression (Figure 5). Additionally, the expression of miR398 targets, *CSDs* and *CCS*, is also reduced in *ago1-25* (Figure 3C).

To test whether the FeSOD1-AGO1 interaction functions reciprocally, we examined the role of FeSOD1 in AGO1 function. FeSOD1 is essential for ROS response, specifically in eliminating superoxide radicals (Ravet and Pilon, 2013). In this sense, the interaction between AGO1 and FeSOD1 suggests that antioxidant protection is necessary for AGO1 function. To check this, we studied the AGO1 protein content in *fsd1-2* mutants. Homozygous *fsd1-2* seedlings grown under Cu-deficient and Cu-excess conditions exhibit reduced ROS neutralization, affecting phenotypes like increased chlorosis (Melicher *et al*., 2022). Moreover, this capacity depends on Cu availability, as *FSD1* expression is regulated by Cu status. AGO1 levels depend on Cu status and the presence of FeSOD1, supporting the hypothesis that Cu deficiency and FeSOD1 have a protective role against oxidative damage to AGO1 (Figure 3B). Furthermore, the effects on different Cu-miRNA targets (*ARPN*, *CCS*, *CSD2*, *GSTU25*, *LAC3*) (Figure 5A) highlights the defect in AGO1 function in the absence of this protective barrier. Taken together, our findings reveal the challenges plants face under Cu deficiency in maintaining RNA interference efficiency, given the essential role of Cu-miRNAs and the stability of AGO1 protein under these conditions. The substitution of Cu/ZnSOD with FeSOD1 serves as a solution to this compromise (Figure 6). Increased Fe demand for FeSOD1 highlights the interaction between Cu and Fe homeostasis (Perea-García *et al*., 2021; Waters *et al*., 2012). Notably, IRT1, the main Fe transporter, was enriched in TST-AGO1 complexes purified under Cu deficiency. Further studies are needed to clarify the role of the AGO1-IRT1 interaction, although an indirect association via IRT1-RNA binding proteins with the RISC may be plausible.

While stress conditions often reveal molecular imbalances, such as RNA interference functioning with unstable AGO1, these balances could also exist under normal conditions, as observed here for AGO1 under Cu sufficiency. Despite low *FSD1* levels under Cu sufficiency, the expression of non-Cu-miRNAs, such as miR167 (*ARF8*) (Wu *et al*., 2006), miR169 (*NF-YA9*) (Sorin *et al*., 2014), miR156 (*SPL3* and *SPL9*) (Kim *et al*., 2012) and miR159 (*TCP4*) (Palatnik *et al*., 2007) was also affected in *fsd1-2* (Figure 5B). These findings suggest a broader scenario (Figure 6) where maintaining an antioxidant environment around AGO1 is crucial for proper RNA interference function, particularly under Cu deficiency.

## MATERIALS AND METHODS

### Plant materials, growth conditions and plant phenotyping

*ago1-25* is a hypomorphic mutant obtained in an EMS mutagenesis and described before (Morel *et al*., 2002). *fsd1-2* (GABI_740E11) is a mutant line derived from a T-DNA insertion and described before (Melicher *et al*., 2022). Wild-type Arabidopsis cv Columbia 0 (Col-0), *ago1-25* and *fsd1-2* seeds were grown on plates where Cu concentration was 1 mM, considered as Cu sufficiency, as previously described (Perea-García *et al*., 2020). For severe Cu deficiency condition, 100 μM of (BCS) was added to the growth medium. Seedlings were grown for 10 days at neutral photoperiodic conditions (12 h light, 23°C/12 h darkness, 16°C). Root length of 10-day old seedlings was measured using ImageJ (Abramoff *et al*., 2004).

### Transgene constructs

*pENTR-pAGO1-3xHA-AGO1* (Carbonell *et al*., 2012) was digested with *Hind*III and *Kpn*I, and the two fragments (7812 bp and 1425 bp) were gel-purified. The 1425 bp fragment was ligated into *Hind*III/*Kpn*I-digested *pBSIIKS*+ to generate *pBS-pAGO1-3xHA*. A 4266 bp linear *pBS-pAGO1* fragment lacking the 3xHA tag was amplified using *pBS-pAGO1-3xHA* as template and oligonucleotide pair D2475/D2476, gel purified and assembled with a 136 bp *TST-AGO1* fragment using a plasmid containing a *TST-GFP* cassette and oligonucleotide pair D2477/2478 in the presence of GeneArt Gibson Assembly HiFi Master Mix (Invitrogen) to generate *pBS-pAGO1-TST*. *pBS-pAGO1-TST* was digested with *Hind*III-*Kpn*I, and the resulting 1437 bp fragment was ligated to the 7812 bp mentioned before, to generate *pENTR-pAGO1-TST-AGO1*. Finally, the *pAGO1-TST-AGO1* cassette was cloned by LR recombination into *pMDC99*, a Gateway-compatible plant transformation vector (Curtis and Grossniklaus, 2003), to generate *pMDC99-pAGO1-TST-AGO1*. Nucleotide and amino acid sequences of TST-AGO1 and TST- GFP are listed in Text S1.

A 6003 bp and an 863 bp fragment was amplified from *pENTR-pAGO1-3xHA-AGO1* (Carbonell *et al*., 2012) and *TST-GFP* plasmid with oligonucleotide pair AC-22/AC-23 and AC-20/AC-21, respectively. Both fragments were assembled in the presence of GeneArt Gibson Assembly HiFi Master Mix (Invitrogen) to generate *pENTR-pAGO1-TST-GFP*. The *pAGO1-TST-GFP* was transferred by LR recombination into *pMDC99* to generate *pMDC99- pAGO1-TST-GFP*.

### RNA and protein preparation

Total RNA form Arabidopsis seedlings or inflorescences was isolated as before (Cisneros *et al*., 2023). Triplicate samples from pools of Arabidopsis seedlings or inflorescences were analyzed. Total protein extracts were prepared in 2X PDB buffer (0.0625 M Tris pH 6.8, 2% SDS, 10% glycerol, 10% mercaptoethanol, 0.02% bromophenol blue) in a 1:5 tissue:buffer ratio.

### Real-time RT-qPCR

cDNA synthesis and qPCR analysis were performed essentially as before (Cisneros *et al*., 2025). Briefly, five hundred ng of DNaseI-treated total RNA was used to produce cDNA from 10-day-old Arabidopsis seedlings with the PrimeScript RT Reagent Kit (Perfect Real Time, Takara) according to manufacturer’s instructions. RT-qPCR was done on optical 96-well plates in a QuantStudio 3 Real-Time PCR system (Thermo Fisher Scientific, Waltham, MA, USA) following these conditions: 20 s at 95°C, followed by 40 cycles of 95°C for 3 s and 60°C for 30 s, with an additional melt curve stage consisting of 15 s at 95°C, 1 min at 60°C and 15 s at 95°C. The 20-μl reaction included 10 μl of 2× TB Green Premix Ex Taq (Takara), 2 μl of diluted complementary DNA (1:5), 0.4 μl of ROX II Reference Dye (50X) and 300 nM of each gene-specific primer. Oligonucleotides used for RT-qPCR are listed in Table S1. Target mRNA expression levels were calculated relative to reference genes *ACT2* and *UBQ10*, using the delta delta cycle threshold comparative method of QuantStudio Design and Analysis software, version 1.5.1 (Thermo Fisher Scientific). Three independent biological replicates, and two technical replicates for each biological replicate were analyzed.

### Small RNA blot assays

Small RNA blot assays and band quantification from radioactive membranes were done as described (Cisneros *et al*., 2022). Oligonucleotides used as probes for sRNA blots are listed in Table S1.

### Protein blot assays

Proteins were separated in NuPAGE Novex 4-12% Bis-Tris gels (Invitrogen), transferred to Protran nitrocellulose membranes (Amersham) and detected by chemiluminescence using specific antibodies and SuperSignal West Pico PLUS chemiluminescent substrate (ThermoFisher Scientific) as before (Cisneros *et al*., 2025). Here, for detection of TST-tagged proteins, StrepMAB-Classic HRP Twin-Strep-Tag antibody (IBA) at a 1:10000 dilution was used. For detection of endogenous AGO1 and HISTONE3 (H3), anti-AGO1 (Agrisera) and anti-Histone 3 (Abcam) were used at 1:10000 and 1:4000 dilutions, respectively, and conjugated with 1:20000 of goat anti-rabbit IgG horseradish peroxidase secondary antibody (ThermoFisher Scientific). Images were acquired with an ImageQuant 800 CCD imager (Cytiva) and analyzed with ImageQuantTL v10.2 (Cytiva). Ponceau red S solution (Thermo Fisher Scientific) staining of membranes was used to verify the global protein content of the samples.

### Statistical analysis

Statistical tests are described in the figure legends. Significant differences were determined with two-tailed Student’s *t-*test (*P* < 0.05).

### Affinity purification of protein complexes

Protein complexes containing TST-tagged protein products were purified from 1.5 g of 12-day-old seedlings by affinity chromatography in native conditions using a 1-ml Strep-Tactin XT spin column (IBA) as previously described (Martinez *et al*., 2016).

### Protein identification by liquid chromatography coupled with tandem mass spectrometry

Protein preparations were separated by SDS-PAGE using 4% and 10% acrylamide for stacking and resolving gel, respectively. Electrophoresis was stopped when the front entered 1 cm into the resolving gel, and the gel was stained with Coomassie. The bands were cut for subsequent in-gel digestion with trypsin (12.5 ng/ul). The digest was acidified with 0.5% TFA to stop digestion. The peptides were desalted using C18 columns according to the manufacturer’s protocol (Pierce C18 Spin Columns, Thermo Scientific). The peptides were dried in a speedvac and resuspended in 15 µL of mobile phase (2% ACN, 0.05% TFA) for injection into HPLC. The mixture of tryptic peptides was analyzed by high-resolution mass spectrometry using an Orbitrap Fusion (Thermo Scientific) in “Data Dependent Acquisition” mode, equipped with a nanoESi ion source. The peptide extract, diluted in 2% acetonitrile (ACN) and 0.05% trifluoroacetic acid (TFA), was first separated in an Ultimate 3000 Dionex UHPLC (Thermo Scientific) under the following chromatographic conditions: preconcentration on a C18 pre-column (PepMap100, 5 µm, 300 µm x 5 mm, Thermo Scientific) at a flow rate of 5 µL/min for 3 minutes in 98% acetonitrile and 0.1% TFA. The chromatographic gradient of 4–40% acetonitrile was applied at a flow rate of 300 nL/min on a C18 nano-column (Acclaim PepMap RSLC 75 µm x 50 cm, Thermo Scientific) over 60 minutes. The total chromatography time was 85 minutes. The Orbitrap Fusion operated in positive mode with a voltage of 2 kV. The mass range in “full scan” mode was 400–1500 m/z in the Orbitrap at a resolution of 120,000, with an AGC target of 4×10⁵ and a dynamic “injection time.” Peptide analysis was performed in “data dependent acquisition” mode, operating in MS/MS Top 50 mode, fragmenting as many high-intensity ions as possible per “full scan” within a cycle time of 3 seconds. Fragmentation spectra were acquired in the linear trap by CID fragmentation at a collision energy of 35%, with a dynamic injection time and an AGC target of 1×10². A dynamic exclusion time of 15 seconds was included for each ion to avoid repetitive ion fragmentation. Protein identification was done using Proteome Discoverer 2.1 (PD2.1, Thermo Fisher Scientific). Proteins were identified with the following conditions: database, Uniprot_Arabidopsis-thaliana + AGO-sequences; tolerance, 10 ppm for Orbitrap and 0.02 Da for Ion Trap; maximum of 1 missed cleavages; *b-ions weight 1 (CID) and 0.3 (HCD); *y-ions weigh: 1 (CID-HCD); dynamic modifications, oxidation (Met); fixed modifications, carbamidomethylation (Cys) (*depends on the fragmentation type: CID or HCD); acceptance criteria, Percolator (Delta CN<0.05); FDR of 1% against decoy database. Resulting Excel sheets contain the identifications at the peptide and protein levels (Data S1). The data included in the report are filtered by a 1% FDR based on the probability values obtained by Percolator. The minimum number of peptides per protein is 1.

### Computational analysis

A previously developed bioinformatic pipeline (Ambrós *et al*., 2021)was used to process all information obtained via proteomics. From the raw proteomic lists, all plant proteins were identified and mapped to Arabidopsis genes (proteins), keeping the normalized MS peak area as a quantitative measure. Specifically, we identified proteins from green plants by GenInfo Identifier (GI) through Mascot analysis and conducted sequence alignments using BLAST to identify individual genes in *A. thaliana*, using data from The Arabidopsis Information Resource (TAIR) version 10 (Lamesch *et al*., 2012). For each hit, the associated amino acid sequence was retrieved from the NCBI database in FASTA format using the Biopython Entrez module. Only alignments with e-values < 0.0001 were considered, and any redundant elements were excluded. The protein list obtained from the Cu deficiency condition was compared against the list obtained from the Cu sufficiency condition in a quantitative manner (*i.e.*, by subtracting the MS peak areas) for each transgenic line, discarding those proteins present in control *pAGO1-TST-GFP* lists for each condition. After the control filtering, if a protein was in one list but not in the other, we considered |log2 fold| = 100 (arbitrary value, denoted by Inf in the figures); positive for proteins detected in Cu deficiency but not in sufficiency and negative for proteins detected in Cu sufficiency but not in deficiency. All genes (proteins) in the lists were functionally annotated according to the information from TAIR10.

### Gene and virus identifiers

Arabidopsis gene identifiers are: *ACT2* (AT3G18780), *AGO1* (AT1G48410), *ARF8* (AT5G37020), *ARPN* (AT2G02850), *CCS* (AT1G12520), *CSD1* (AT1G08830), *CSD2* (AT2G28190), *FSD1* (AT4G25100), *GSTU25* (AT1G17180), *LAC3* (AT2G30210), *MSD1* (AT3G10920) *NF-YA9* (AT3G20910), *SPL3* (AT2G33810), *SPL9* (AT2G42200), *TCP4* (AT3G15030), *UBQ10* (AT4G05320).

## ACKOWLEDGEMENTS

This work was supported by grants or fellowships from MCIN/AEI/10.13039/501100011033 (co-financed by the European Regional Development Fund) [RTI2018-095118-A-I00, PID2021-122186OB-I00 and RYC-2017-21648 to A.C.; PID2021-127671NB-I00 to G.R.; PID2020-116940RB-I00 to L.P. and A.P.G.; BIO2017-83184-R and PID2020-114691RB-I00 to J.A.D.; PRE2019-088439 to A.E.C.], and from the Generalitat Valenciana (CIPROM/2022/21 to G.R. and J.A.D.) We thank Dr. Tomás Takác for sharing *fsd1-2* seeds and Carlos Fuentes from the Proteomics Unit of University of Córdoba for the assistance in preparing and analyzing purified proteins.

## CONFLICTS OF INTEREST

The authors declare no conflict of interest.

## AUTHOR CONTRIBUTIONS

A.C. and L.P. planned and designed the research. A.H.T., A.P.G., A.C., A.E.C., L.L.D., A.A.B. and M.O. performed most of the experiments. J.S.V. and J.A.D. performed AGO1 affinity purifications. G.R. did the computational analysis of AP-MS samples. A.H.T., A.P.G, L.P and A.C. analyzed data. A.C. and L.P. wrote the manuscript with input from rest of the authors.

## DATA AVAILABILITY STATEMENT

All data relating to this manuscript can be found within the manuscript and its supplementary files. Data that support the findings of this study are available from the corresponding authors upon reasonable request. The mass spectrometry proteomics data have been deposited to the ProteomeXchange Consortium via the PRIDE (Perez-Riverol *et al*., 2022) partner repository with the dataset identifier PXD057108.

## SUPPLEMENTAL DATA

**Data S1**. Proteins identified by AP-MS in *pAGO1:TST-AGO1* and *pAGO1:TST-GFP* transgenic lines grown under Cu deficiency and Cu sufficiency.

**Data S2.** Proteins identified by AP-MS in each of the three *pAGO1:TST-AGO1* transgenic lines grown under Cu deficiency versus Cu sufficiency, and not present in *pAGO1:TST-GFP* AP-MS samples for each condition.

**Data S3.** Proteins identified by AP-MS in all three *pAGO1:TST-AGO1* transgenic lines grown under Cu deficiency versus Cu sufficiency, and not present in *pAGO1:TST-GFP* AP-MS samples for each condition.

**Figure S1.** Accumulation of miRNA target mRNAs in different Arabidopsis genotypes under Cu deficiency.

**Table S1.** Name, sequence and use of oligonucleotides used in the present study.

**Text S1**. Nucleotide and amino acid sequences of TST-based inserts.

## Notes

### Competing Interest Statement

The authors have declared no competing interest.

### Summary of Updates

This version of the manuscript has been revised to inbclude the supplementary material.

