## Supplemental figure-table-text for "*Arabidopsis thaliana* Iron Superoxide Dismutase FeSOD1 Protects ARGONAUTE 1 in a Copper-Dependent Manner"

### SUPPLEMENTARY INFORMATION

**Data S1**. Proteins identified by AP-MS in *pAGO1:TST-AGO1* and *pAGO1:TST-GFP* transgenic lines grown under Cu deficiency and Cu sufficiency.

**Data S2.** Proteins identified by AP-MS in each of the three *pAGO1:TST-AGO1* transgenic lines grown under Cu deficiency versus Cu sufficiency, and not present in *pAGO1:TST-GFP* AP-MS samples for each condition.

**Data S3.** Proteins identified by AP-MS in all three *pAGO1:TST-AGO1* transgenic lines grown under Cu deficiency versus Cu sufficiency, and not present in *pAGO1:TST-GFP* AP-MS samples for each condition.

**Figure S1**. Accumulation of miRNA target mRNAs in different Arabidopsis genotypes under Cu deficiency.

**Table S1.** Name, sequence and use of oligonucleotides used in the present study.

**Text S1**. Nucleotide and amino acid sequences of TST-based inserts.

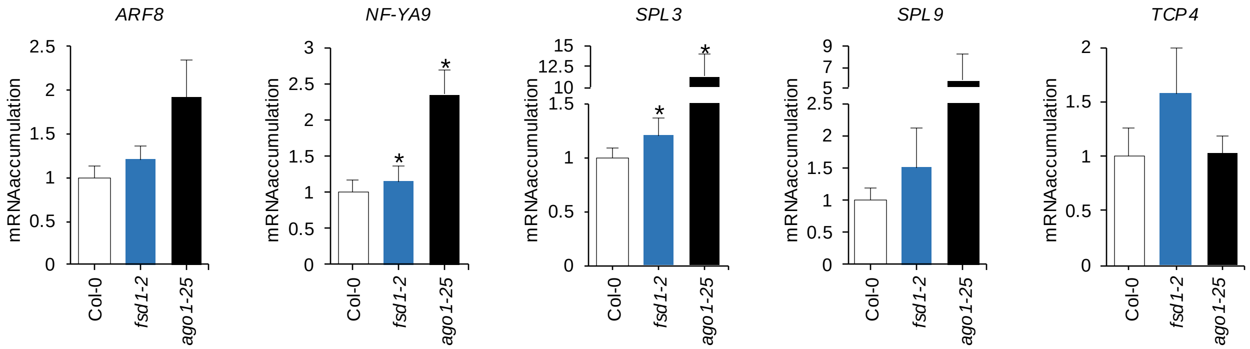


**Figure S1 . Accumulation of miRNA target mRNAs in different *Arabidopsis thaliana* genotypes under Cu deficiency.** The relative expression of *ARF8*, *NF-YA9*, *SPL3*, *SPL9* and *TCP4* was determined by RT-qPCR in 10-day-old Col-0, *fsd1-2* and *ago1-25* seedlings, grown under Cu deficiency (100 µM BCS). Other details are as in Figure 3A and 3C.

**Table S1.** Name, sequence and use of DNA oligonucleotides used in this study.

| Name | Sequence | Type* | Construct/Aim |
|---|---|---|---|
| AC-20 | cggccgccccccttcaccatgAGCGCATGGAGTCATCCTC | ssDNA | Generation of *pMDC99-pAGO1-TST-GFP*. |
| AC-21 | tcaacggcgcgcccaccccttTTACTTGTACAGCTCGTCCATG | ssDNA | |
| AC-22 | AAGGGTGGGCGCGCCGTT | ssDNA | |
| AC-23 | CATGGTGAAGGGGGGCGGC | ssDNA | |
| AC-55 | AGGGGCCATGCTAATCTTCTC | ssDNA | Probe for U6 detection. |
| AC-159 | AAAAATGGCTGAGGCTGATGA | ssDNA | qPCR amplification of *ACT2* mRNA |
| AC-160 | GAAAAACAGCCCTGGGAGC | ssDNA | |
| AC-457 | T+ACG+CTA+TGT+TGG+ACT+TAG+AA | ssLNA | Probe for tasiR255 detection. |
| AC-1055 | TACCACGAGCTGCGAGAAGAGT | ssDNA | qPCR amplification of *ARF8* mRNA |
| AC-1056 | TTGCGGGAATAACCCACCACTG | ssDNA | |
| AC-1057 | CAAGGTTCAGTTGGTGGAGGA | ssDNA | qPCR amplification of *SPL9* mRNA |
| AC-1058 | TGAAGAAGCTCGCCATGTATTG | ssDNA | |
| AC-1196 | GGGTACTGGAGGACGGTTTG | ssDNA | qPCR amplification of *NF-YA9* mRNA |
| AC-1197 | CGGGGACTGAGTAACATGACC | ssDNA | |
| AC-1200 | CAACCGATACAGGAAACGGAG | ssDNA | qPCR amplification of *TCP4* mRNA |
| AC-1201 | CTGGTATGCGAAAACCCGAAG | ssDNA | |
| AC-1290 | TCGGTGGACAGAAGTGGGAATA | ssDNA | qPCR amplification of *AGO1* mRNA |
| AC-1291 | CCAAAGAACGTGGTAATGAGCAGG | ssDNA | |
| ARPN-F | TGACTCTCATGGCTGTGTCA | ssDNA | qPCR amplification of *ARPN* mRNA |
| ARPN-R | CACTACGTTGTGCATCCTCG | ssDNA | |
| CCS-F | TCTCCACGTCTCTTGGGACTTT | ssDNA | qPCR amplification of *CCS* mRNA |
| CCS-R | AGCTGAGGCATGGCTCGAT | ssDNA | |
| CSD1-F | CATCATTGGTCTCCAGGGCT | ssDNA | qPCR amplification of *CSD1* mRNA |
| CSD1-R | GACCTCCTTATTACATCAAT | ssDNA | |
| CSD2-F | GTCCTACAACTGTGAAT | ssDNA | qPCR amplification of *CSD2* mRNA |
| CSD2-R | TCCATGAGGCCCTGGAGT | ssDNA | |
| D2475 | CATGGTGAAGGGGGGCGGC | ssDNA | Generation of *pMDC99-pAGO1-TST-AGO1* |
| D2476 | GTGAGAAAGAGAAGAACGG | ssDNA | |
| D2477 | cggccgccccccttcaccatgAGCGCATGGAGTCATCCTC | ssDNA | |
| D2478 | tccgttcttctctttctcacTCCGGATTTTTCGAACTG | ssDNA | |
| FSD1-F | ACCGAAGACCAGATTACATA | ssDNA | qPCR amplification of *FSD1* mRNA |
| FSD1-R | TGGCACTTACAGCTTCCCAA | ssDNA | |
| GSTU25-F | TGGCAGACGAGGTGATTCTT | ssDNA | qPCR amplification of *GSTU25* mRNA |
| GSTU25-R | TTGCTAGGCCAAACTTCGTC | ssDNA | |
| LAC3-F | AACTGCTTTCACCAACCGTC | ssDNA | qPCR amplification of *LAC3* mRNA |
| LAC3-R | TGGTAGCACGAAGGACATGT | ssDNA | |
| MSD1-F | GAAGAAGCTAGTTGTTGACAC | ssDNA | qPCR amplification of *MSD1* mRNA |
| MSD1-R | CCTCGCTTGCATATTTCCAG | ssDNA | |
| UBQ10-F | TAATCCCTGATGAATAAGTGTTCTAC | ssDNA | qPCR amplification of *ACT2* mRNA |
| UBQ10-R | AAAACGAAGCGATGATAAAGAAG | ssDNA | |

*ssDNA: single-stranded DNA; dsDNA: double-stranded DNA; LNA: locked nucleic acid.

**Text S1.** Nucleotide and amino acid sequences of TST-based inserts.

1. Nucleotide sequences

>TST-GFP
AGCGCATGGAGTCATCCTCAATTCGAGAAAGGTGGAGGTTCTGGCGGTGGATCGGGAGGTTCAGCGTGGAGCCAC
CCGCAGTTCGAAAAATCCGGAATGGTGAGCAAGGGCGAGGAGCTGTTCACCGGGGTGGTGCCCATCCTGGTCGAG
CTGGACGGCGACGTAAACGGCCACAAGTTCAGCGTGTCCGGCGAGGGCGAGGGCGATGCCACCTACGGCAAGCTG
ACCCTGAAGTTCATCTGCACCACCGGCAAGCTGCCCGTGCCCTGGCCCACCCTCGTGACCACCCTGACCTACGGC
GTGCAGTGCTTCAGCCGCTACCCCGACCACATGAAGCAGCACGACTTCTTCAAGTCCGCCATGCCCGAAGGCTAC
GTCCAGGAGCGCACCATCTTCTTCAAGGACGACGGCAACTACAAGACCCGCGCCGAGGTGAAGTTCGAGGGCGAC
ACCCTGGTGAACCGCATCGAGCTGAAGGGCATCGACTTCAAGGAGGACGGCAACATCCTGGGGCACAAGCTGGAG
TACAACTACAACAGCCACAACGTCTATATCATGGCCGACAAGCAGAAGAACGGCATCAAGGTGAACTTCAAGATC
CGCCACAACATCGAGGACGGCAGCGTGCAGCTCGCCGACCACTACCAGCAGAACACCCCCATCGGCGACGGCCCC
GTGCTGCTGCCCGACAACCACTACCTGAGCACCCAGTCCGCCCTGAGCAAAGACCCCAACGAGAAGCGCGATCAC
ATGGTCCTGCTGGAGTTCGTGACCGCCGCCGGGATCACTCTCGGCATGGACGAGCTGTACAAGTAA
TST
GFP
ATG Native GFP start codon.
TAA Added STOP codon.

>TST-AGO1
AGCGCATGGAGTCATCCTCAATTCGAGAAAGGTGGAGGTTCTGGCGGTGGATCGGGAGGTTCAGCGTGGAGCCAC
CCGCAGTTCGAAAAATCCGGAGTGAGAAAGAGAAGAACGGATGCTCCATCTGAAGGAGGTGAAGGCTCTGGGTCT
CGTGAAGCTGGTCCAGTCTCAGGTGGTGGACGTGGTTCACAGCGAGGTGGTTTCCAGCAGGGAGGAGGACAACAC
CAAGGTGGAAGGGGTTATACTCCTCAACCTCAACAGGGAGGTCGTGGTGGTCGTGGATATGGGCAACCACCACAA
CAGCAACAACAGTATGGAGGACCACAAGAGTACCAAGGAAGAGGAAGAGGAGGACCTCCTCATCAAGGAGGTCGA
GGAGGGTATGGCGGTGGCCGTGGAGGTGGACCTTCTTCTGGACCACCGCAGAGACAATCAGTTCCCGAGCTGCAT
CAAGCTACCTCACCTACTTATCAAGCGGTGTCTTCTCAGCCTACACTGTCTGAGGTGAGTCCTACCCAGGTACCA
GAACCTACTGTTCTGGCTCAGCAATTTGAACAACTCTCTGTTGAACAAGGAGCTCCCAGTCAGGCAATCCAGCCT
ATACCTTCTTCTAGCAAGGCTTTCAAGTTTCCAATGAGGCCTGGTAAAGGACAGAGTGGAAAGCGTTGCATTGTG
AAGGCTAACCATTTCTTTGCTGAACTGCCTGATAAGGATTTGCACCATTATGATGTTACCATTACTCCGGAAGTT
ACATCAAGGGGTGTCAATCGTGCTGTGATGAAACAACTTGTTGATAATTATCGTGATTCTCACCTTGGAAGTCGT
CTTCCAGCGTATGATGGTCGAAAAAGTCTTTACACTGCTGGTCCACTTCCCTTTAACTCCAAGGAGTTCAGAATC
AATCTTCTTGACGAAGAAGTAGGGGCTGGAGGTCAAAGACGAGAAAGGGAATTTAAAGTTGTGATCAAGCTAGTT
GCACGTGCTGATCTGCATCACCTAGGAATGTTTTTGGAGGGGAAACAATCAGATGCCCCACAGGAAGCTCTGCAG
GTTCTTGACATTGTTCTTCGTGAGCTGCCGACCTCTAGAATCAGGTATATTCCGGTGGGCCGGTCCTTTTATTCC
CCTGATATAGGAAAAAAACAATCATTGGGGGATGGCTTGGAGAGCTGGCGTGGATTCTACCAAAGCATTCGTCCT
ACACAGATGGGCTTATCACTCAATATTGATATGTCATCGACAGCCTTCATAGAGGCAAACCCTGTGATTCAGTTT
GTCTGTGATTTGCTTAACCGGGATATTTCTTCTCGACCTTTATCTGATGCTGATCGTGTTAAGATAAAAAAGGCT
CTTAGAGGTGTCAAAGTTGAAGTGACTCATCGAGGAAACATGCGCCGGAAGTACCGCATTTCCGGTTTGACTGCT
GTGGCCACTCGGGAATTGACATTCCCAGTAGATGAAAGAAATACTCAGAAATCTGTTGTAGAATACTTCCACGAA
ACATATGGTTTTCGCATTCAGCACACTCAACTACCATGCTTGCAAGTTGGGAATTCTAATAGGCCTAATTACTTA
CCAATGGAGGTATGCAAGATTGTTGAAGGCCAGCGGTATTCCAAAAGATTGAATGAGAGACAGATCACTGCTTTG
CTGAAGGTTACCTGTCAGCGCCCGATAGATCGAGAAAAAGATATCTTACAGACGGTGCAACTCAATGATTATGCT
AAAGATAATTATGCTCAAGAGTTTGGCATCAAAATAAGTACTTCTCTGGCTTCTGTTGAGGCTCGTATACTGCCT
CCTCCATGGCTTAAGTACCACGAGTCTGGAAGGGAAGGGACTTGTCTGCCACAAGTTGGTCAATGGAACATGATG
AATAAGAAAATGATCAATGGTGGAACGGTGAATAATTGGATCTGCATCAACTTTTCTAGGCAAGTGCAGGACAAT
CTAGCGCGTACATTTTGTCAGGAACTTGCTCAAATGTGTTACGTATCTGGCATGGCATTTAATCCGGAACCAGTC
CTCCCACCAGTCAGTGCTCGCCCTGAGCAAGTAGAGAAGGTCTTGAAGACTAGATATCATGATGCCACATCAAAA
CTCTCCCAAGGAAAAGAAATTGATCTGCTTATTGTCATTCTGCCCGATAATAATGGATCATTATACGGTGATTTG
AAACGCATATGTGAGACTGAACTCGGCATAGTCTCTCAATGTTGCCTGACAAAGCATGTCTTTAAGATGAGCAAA
CAATACATGGCTAATGTTGCGCTGAAGATTAATGTGAAGGTTGGAGGAAGAAACACAGTGCTTGTTGATGCTCTA
TCTAGGCGGATTCCTCTAGTCAGTGATCGACCCACCATTATATTTGGTGCTGATGTTACCCACCCTCACCCTGGA
GAGGATTCAAGCCCATCTATTGCTGCTGTTGTGGCATCTCAGGATTGGCCTGAAATCACTAAATATGCTGGATTA
GTTTGCGCTCAAGCGCATAGGCAGGAGCTCATTCAGGATCTGTTCAAAGAGTGGAAGGATCCTCAGAAAGGTGTG
GTGACTGGTGGCATGATAAAGGAGTTGCTCATAGCCTTCCGTAGATCAACTGGGCATAAACCACTAAGGATCATC
TTCTACAGGGATGGAGTCAGTGAGGGACAATTTTACCAAGTTTTGCTCTATGAACTTGATGCCATCCGCAAGGCC
TGTGCTTCGCTGGAAGCAGGTTATCAACCACCAGTGACATTTGTGGTGGTGCAGAAGCGTCATCACACGAGGCTG
TTTGCTCAGAACCACAATGATCGCCATTCGGTGGACAGAAGTGGGAATATTTTACCTGGCACTGTTGTGGACTCT

**AAAATCTGCCACCCTACAGAGTTTGACTTTTACCTCTGTAGTCATGCTGGTATTCAGGGCACTTCTCGACCTGCT
CATTACCACGTTCTTTGGGATGAGAACAACTTTACTGCAGATGGACTTCAATCTCTGACCAATAACTTATGTTAC
ACGTATGCAAGATGCACACGCTCAGTTTCAATTGTTCCCCCTGCATATTATGCACATCTAGCAGCTTTTAGGGCT
CGATTCTACATGGAGCCAGAGACATCAGACAGTGGCTCAATGGCTAGTGGGAGCATGGCACGTGGAGGTGGAATG
GCTGGTAGAAGCACACGCGGGCCTAATGTCAATGCTGCAGTGAGGCCACTCCCAGCTCTGAAAGAGAATGTGAAG
CGTGTCATGTTCTACTGCTGA**

**TST**

**AGO1**

### 2. Amino acid sequences

>TST-GFP

**SAWSHPQFEKGGGSGGGSGGSAWSHPQFEKSGMVSKGEELFTGVVPILVELDGDVNGHKFSVSGEGEGDATYGKL
TLKFICTTGKLPVPWPTLVTTLTYGVQCFSRYPDHMKQHDFFKSAMPEGYVQERTIFFKDDGNYKTRAEVKFEGD
TLVNRIELKGIDFKEDGNILGHKLEYNYNSHNVYIMADKQKNGIKVNFKIRHNIEDGSVQLADHYQQNTPIGDGP
VLLPDNHYLSTQSALSKDPNEKRDHMVLLEFVTAAGITLGMDELYK**

>TST-AGO1

**SAWSHPQFEKGGGSGGGSGGSAWSHPQFEKSGVRKRRTDAPSEGGEGSGSREAGPVSGGGRGSQRGGFQQGGGQH
QGGRGYTPQPQQGGRGGRGYGQPPQQQQQYGGPQEYQGRGRGGPPHQGGRGGYGGGRGGGPSSGPPQRQSVPELH
QATSPTYQAVSSQPTLSEVSPTQVPEPTVLAQQFEQLSVEQGAPSQAIQPIPSSSKAFKFPMRPGKGQSGKRCIV
KANHFFAELPDKDLHHYDVTITPEVTSRGVNRAVMKQLVDNYRDSHLGSRLPAYDGRKSLYTAGPLPFNSKEFRI
NLLDEEVGAGGQRREREFKVVIKLVARADLHHLGMFLEGKQSDAPQEALQVLDIVLRELPTSRIRYIPVGRSFYS
PDIGKKQSLGDGLESWRGFYQSIRPTQMGLSLNIDMSSTAFIEANPVIQFVCDLLNRDISSRPLSDADRVKIKKA
LRGVKVEVTHRGNMRRKYRISGLTAVATRELTFPVDERNTQKSVVEYFHETYGFRIQHTQLPCLQVGNSNRPNYL
PMEVCKIVEGQRYSKRLNERQITALLKVTCQRPIDREKDILQTVQLNDYAKDNYAQEFGIKISTSLASVEARILP
PPWLKYHESGREGTCLPQVGQWNMMNKKMINGGTVNNWICINFSRQVQDNLARTFCQELAQMCYVSGMAFNPEPV
LPPVSARPEQVEKVLKTRYHDATSKLSQGKEIDLLIVILPDNNGSLYGDLKRICETELGIVSQCCLTKHVFKMSK
QYMANVALKINVKVGGRNTVLVDALSRRIPLVSDRPTIIFGADVTHPHPGEDSSPSIAAVVASQDWPEITKYAGL
VCAQAHRQELIQDLFKEWKDPQKGVVTGGMIKELLIAFRRSTGHKPLRIIFYRDGVSEGQFYQVLLYELDAIRKA
CASLEAGYQPPVTFVVVQKRHHTRLFAQNHNDRHSVDRSGNILPGTVVDSKICHPTEFDFYLCSHAGIQGTSRPA
HYHVLWDENNFTADGLQSLTNNLCYTYARCTRSVSIVPPAYYAHLAAFRARFYMEPETSDSGSMASGSMARGGGM
AGRSTRGPNVNAAVRPLPALKENVKRVMFYC**
